# Data driven and cell specific determination of nuclei-associated actin structure

**DOI:** 10.1101/2023.04.06.535937

**Authors:** Nina Nikitina, Nurbanu Bursa, Matthew Goelzer, Madison Goldfeldt, Chase Crandall, Sean Howard, Janet Rubin, Aykut Satici, Gunes Uzer

## Abstract

Quantitative and volumetric assessment of filamentous actin fibers (F-actin) remains challenging due to their interconnected nature, leading researchers to utilize threshold based or qualitative measurement methods with poor reproducibility. Here we introduce a novel machine learning based methodology for accurate quantification and reconstruction of nuclei-associated F-actin. Utilizing a Convolutional Neural Network (CNN), we segment actin filaments and nuclei from 3D confocal microscopy images and then reconstruct each fiber by connecting intersecting contours on cross-sectional slices. This allowed measurement of the total number of actin filaments and individual actin filament length and volume in a reproducible fashion. Focusing on the role of F-actin in supporting nucleocytoskeletal connectivity, we quantified apical F-actin, basal F-actin, and nuclear architecture in mesenchymal stem cells (MSCs) following the disruption of the Linker of Nucleoskeleton and Cytoskeleton (LINC) Complexes. Disabling LINC in mesenchymal stem cells (MSCs) generated F-actin disorganization at the nuclear envelope characterized by shorter length and volume of actin fibers contributing a less elongated nuclear shape. Our findings not only present a new tool for mechanobiology but introduce a novel pipeline for developing realistic computational models based on quantitative measures of F-actin.

## Introduction

Mechanical information is translated into biological response through perturbations of a highly organized and connected F-actin cytoskeleton where LINC-mediated F-actin connections to the nucleus at the “actin cap” translate these mechanical forces into the nucleus to alter both nuclear structure and gene expression^1^. While understanding the organization and connectivity of cytoskeletal networks has been a research topic in cell mechanobiology for many years^2^, reconstructing the interconnected structures of branching F-actin fibers remains a technical barrier. Much of the information regarding F-actin organization relies on manual and semi-automated processing of 2D images through open-source programs such as imageJ^3^; these methods yield qualitative information with poor quantitative numbers for molecular structures. Furthermore, 3D reconstruction of F-actin represents another challenge as these methods generally represent the cytoskeleton as a simple planar geometry^4–6^ precluding the development of models that recapture the full complexity of cellular cytoskeletal networks. To provide quantitative information from planar analysis, fluorescence and electron microscopy-based image analysis methods have been developed to analyze biopolymer properties, including number, length, and organization. Reconstruction methods typically involve enhancing filamentous features to identify and isolate filaments in images, separating filaments from the background, and extracting individual filaments using line segment detectors^7–10^. Aside from these planar approaches, only few studies have been conducted on the process of separating cytoskeletal filaments and networks in 3D using immunofluorescence microscopy images^11, 12^. While such methods have provided valuable information, reproducible volumetric quantification of F-actin is still beyond the reach of many research groups necessitating an easy to use, unbiased and repeatable volumetric reconstruction of dynamic cytoskeletal networks in cells.

LINC complexes formed by a family of proteins that include KASH (Klarsicht, ANC-1, Syne Homology) and SUN (Sad1p, UNC-84) domains provide mechanical connection between cytoplasmic and nuclear compartments. LINC-mediated nucleo-cytoplasmic connectivity has been shown to play important roles in mechanosensitivity,^13, 14^ chromatin organization,^15–17^ and DNA repair mechanisms.^18, 19^ Here, utilizing intact and LINC-disabled MSCs, we describe a novel approach for automating reconstruction of the nucleo-cytoskeletal architecture that is based on deep learning-assisted image analysis and segmentation of cross-sectional image slices. Focused on the actin fiber architecture within the nuclear region, our method provides precise reconstruction of nuclei-associated fibers, nuclei, and enables extraction of associated statistical data. The software package developed for this method, “afilament”, is publicly available for use and validation (see data availability section). Using the afilament software and confocal images of primary MSCs, we have further quantitatively compared the cell specific consequences of disabling LINC complex on the F-actin cytoskeleton.

### Methods and Data Collection

#### Cell Culture

Bone marrow derived MSCs (mdMSC) from 8-10 wk male C57BL/6 mice were isolated as described from multiple mouse donors and MSCs pooled, providing a heterogenous MSCs cell line.^20^ Briefly, tibial and femoral marrow were collected in RPMI-1640, 9% FBS, 9% HS, 100 μg/ml pen/strep and 12μM L-glutamine. After 24 hours, non-adherent cells were removed by washing with phosphate-buffered saline and adherent cells were cultured for 4 weeks. Passage 1 cells were collected after incubation with 0.25% trypsin/1 mM EDTA × 2 minutes and re-plated in a single 175-cm^2^ flask. After 1-2 weeks, passage 2 cells were re-plated at 50 cells/cm^2^ in expansion medium (Iscove’s modified Dulbecco’s Medium (IMDM), 9% FBS, 9% HS, antibiotics, L-glutamine). mdMSCs were re-plated every 1-2 weeks for two consecutive passages up to passage 5 and tested for osteogenic and adipogenic potential, and subsequently frozen.

These isolated MSC stocks were stably transduced with a doxycycline-inducible plasmid expressing an mCherry tagged dominant-negative KASH (dnKASH) domain^21^. The dnKASH plasmid was lentiviral packaged as a generous gift from Dr. Daniel Conway (Addgene # 125554). Lentivius supernatant was added to growth media with polybrene (5 μg/ml). Lentivirus growth media mixture was added to 50-70% confluent MSCs. Lentivirus media was replaced 48 hours later with selection media containing G418 (1mg/ml) for 5 days to select stably infected dnKASH-MSCs. Calf serum (CS) was obtained from Atlanta Biologicals (Atlanta, GA). MSCs were maintained in IMDM with FBS (10%, v/v) and penicillin/streptomycin (100μg/ml). For immunostaining experiments, seeding cell density was 3,000 cells per cm^2^ in growth media. Twenty-four hours after seeding, dnKASH cells were given growth media containing doxycycline (1 μg/ml).

### RNA-seq

Total RNA was extracted using RNAeasy (Qiagen) for three samples per group. Total RNA samples were sent to Novogene for mRNA sequencing and analysis. Briefly, the index of the reference genome was built using Hisat2 v2.0.5 and paired-end clean 2 reads were aligned to the reference genome using Hisat2 v2.0.5. featureCounts v1.5.0-p3 was used to count the reads numbers mapped to each gene. Differential expression analysis was performed using the DESeq2 R package (1.20.0). DESeq2 provides statistical routines for determining differential expressions in digital gene expression data using a model based on the negative binomial distribution. The resulting P-values were adjusted using the Benjamini and Hochberg’s approach for controlling the false discovery rate. Genes with an adjusted p-value < 0.05 and fold-change (FC) > 0.2 found by DESeq2 were assigned as differentially expressed. Genes with significant differential gene expression were further analyzed with DAVID for pathway analysis^22^. Pathways with a p < 0.05 were selected.

### Immunofluorescence

Forty-eight hours after dnKASH expression, cells were fixed with 4% paraformaldehyde. Cells were permeabilized by incubation with 0.1% Triton X-100. Cells were incubated in a blocking serum consisting of PBS with 1% Bovine Serum Albumin. For nuclear staining, cells were incubated with NucBlue Hoechst 33342 stain (Fisher Scientific) according to the manufacturer’s protocol. F-actin was stained using Phalloidin (iFluor 488, Cayman Chemicals). Reagents used for immunofluorescence and their concentrations are listed in **Supplementary Table S1**. The fluorescent actin cytoskeleton images were obtained using a Leica Stellaris 5 confocal system configured with a Leica DMi8 inverted microscope and 63x/1.4 Oil HC PL APO objective.

### Reconstruction of apical and basal actin stress fibers of MSC from confocal microscope images

The reconstruction algorithm is divided into three phases: confocal image preprocessing, deep learning image segmentation, and postprocessing (**Fig. 1**).

**Figure 1:**
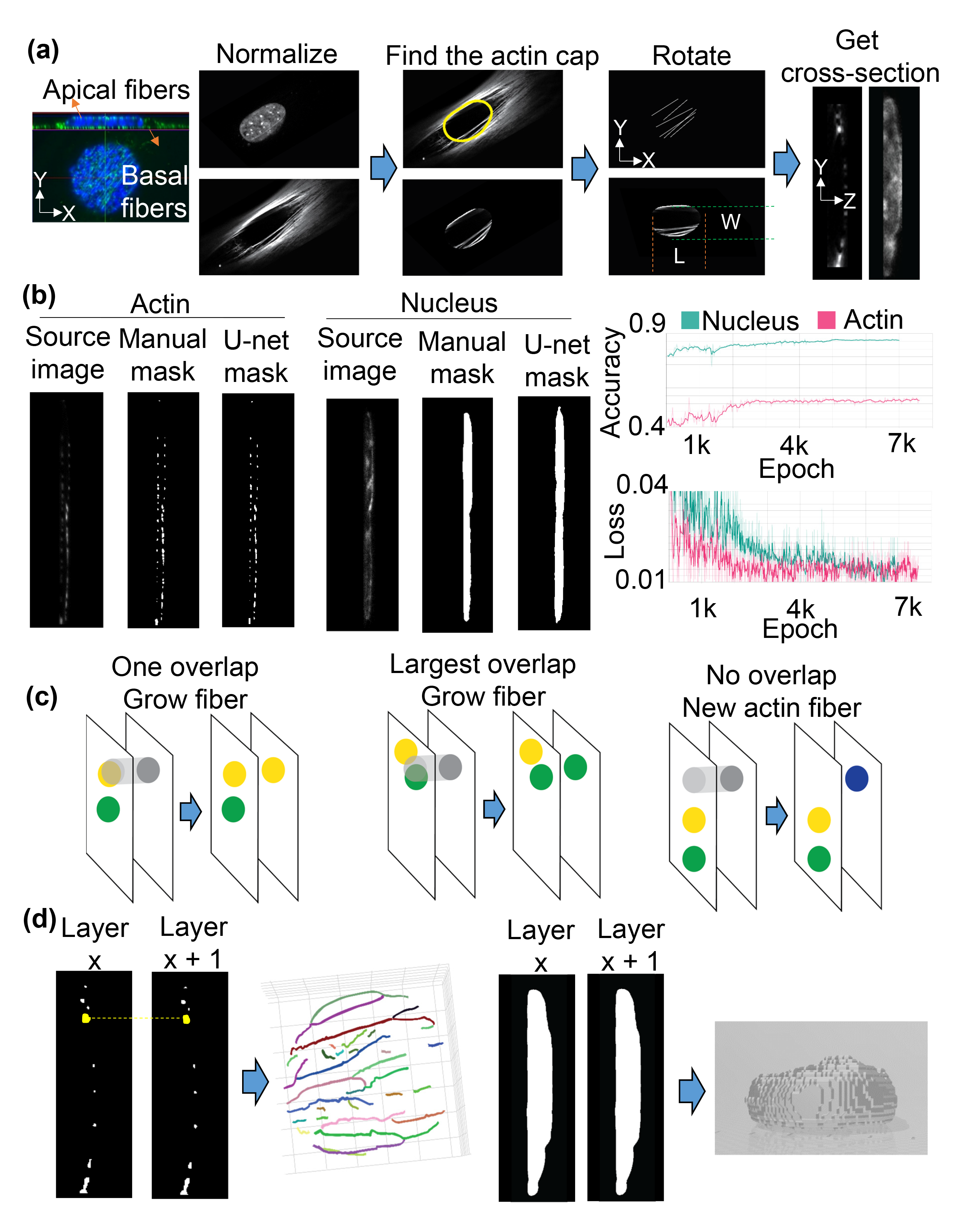
Stress Fiber Reconstruction Algorithm. **(a)** Image preprocessing algorithm reads images and metadata, normalizes the images, cuts out the region corresponding to the nucleus location, rotates the image to align fibers based on Hough Transformation (Methods) of maximal projection of a fiber layer, and converts the processed single-cell image layers into cross-section layers. **(b)** Training details for the neural network. The learning parameters used were a learning rate of 0.001, a batch size of 1, and 200 epochs. The loss function, which quantifies the differences between predicted and target images, was minimized during training. To prioritize false-negative results for actin fibers, the loss function was adjusted to increase the error by 200. **(c)** Reconstruction of actin fibers and nuclei models the initial actin fibers based on the biggest intersection of actin contour on successive layers of actin masks and **(d)** reconstructs nuclei combining contour of nuclei cross-section masks layer by layer.

**Figure 2:**
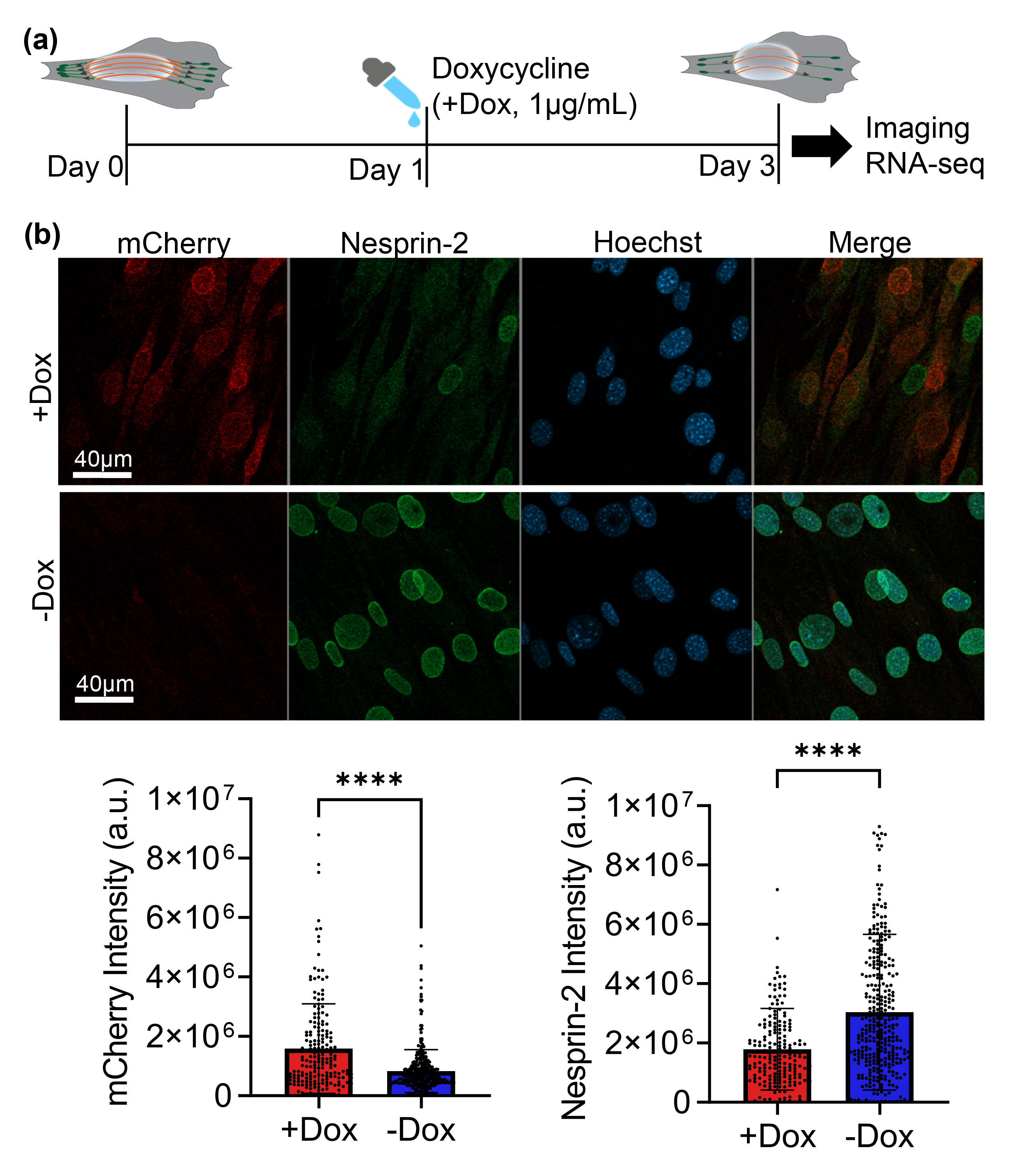
Overexpressing Nesprin KASH domain disables LINC function in MSCs. **(a)** KASH expression was induced in MSCs harboring a doxycycline (Dox) inducible mCherry-tagged KASH domain by adding 1 μg/ml Dox to cell culture medium. No Dox treatment was used as control. Imaging and RNAseq outcomes were acquired 48 hours after the Dox treatment at day 3 after cell seeding. **(b)** +Dox treatment resulted increased mCherry intensity by 73% (n=515, p<0.0001) and decreased Nesprin-2 intensity by 63% (n = 530, P < 0.0001).

**Figure 3:**
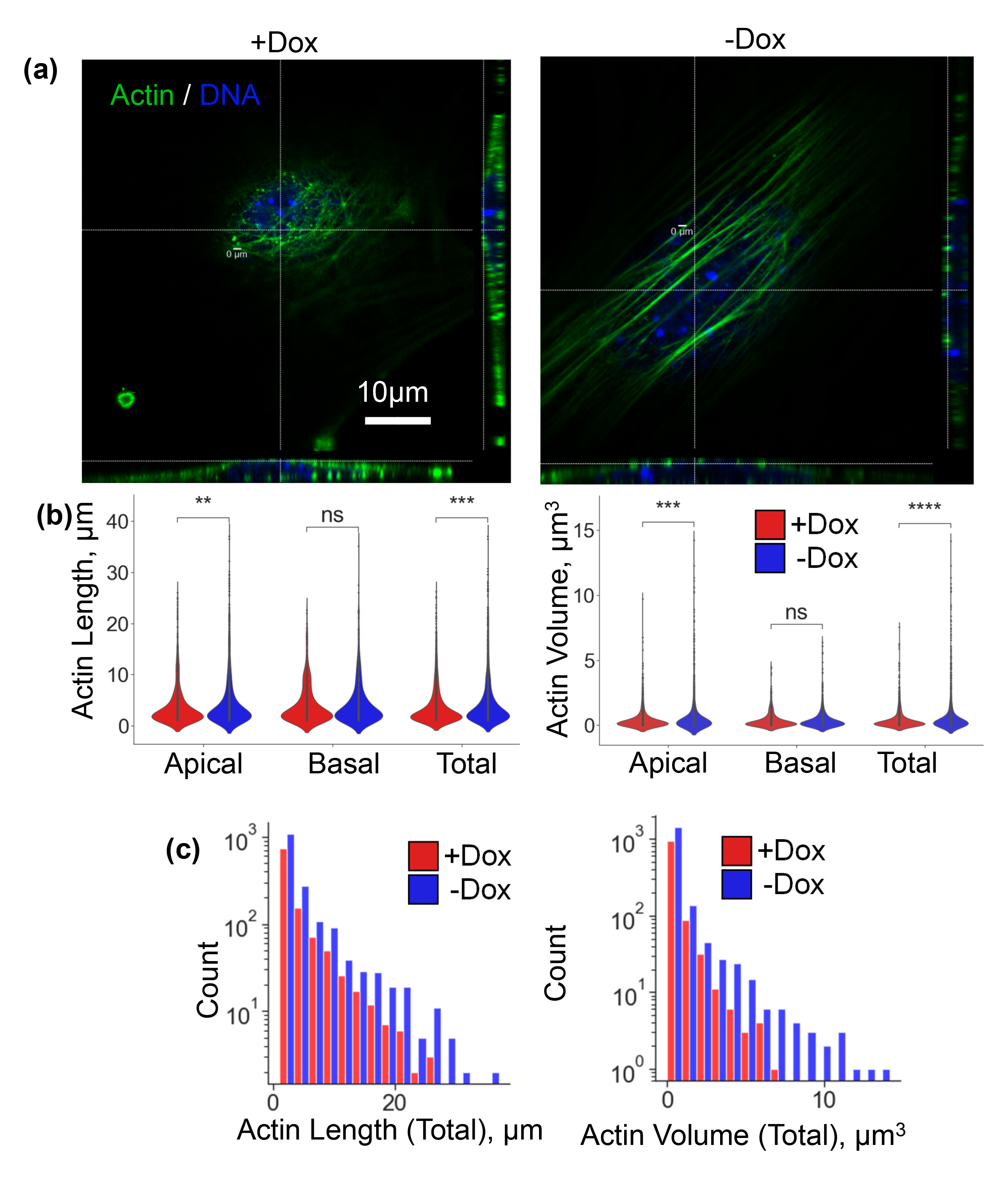
Disabling LINC function reduces unbiased measures of apical but not basal f- actin. **(a)** Visualization of +Dox treatment indicates less and disorganized f-actin fibers across the apical nuclear surface. **(b)** When fibers from all cells combined, +Dox treatment resulted in shortened total and apical fiber lengths (15%, p<0.001). Total fiber volume and apical fiber volume were 30% (p<0.001) and 32% (p<0.001) smaller in the +Dox groups, respectively. No changes in basal actin were observed. **(c)** Distribution of total actin length and volume showed that -Dox treatment exhibited longer actin fibers with larger volumes.

#### Preprocessing

The preprocessing phase (**Fig. 1a**) prepares images for segmentation. The algorithm acquires image resolution, z-stack size and bit depth via the python-bioformats package^23^. To detect nuclear area, thresholding was applied to each layer of the confocal image z-stack to isolate the largest circumference, which was then cropped for further analysis. To align F-actin fibers along the nucleus, fiber layers were converted into 2D via maximal projection and gross fiber orientation was detected via Hough Transform and aligned along the X axis. This rotation allows detection of fibers through Y-Z cross sectional view along the X-axis. As apical and basal fibers are not necessarily parallel to each other^24^, we preprocessed basal and apical fibers separately. To optimize reconstruction efficiency for apical fiber analysis, rotation was based on maximal projection of the top 50% of the height and the bottom 50% was designated as the basal fibers. Finally, these rotated sets of nuclei and F-actin images were converted into Y-Z cross-sectional layer-sets along the X axis.

#### Image segmentation

Shown in **Fig.1b**, actin fibers on Y-Z cross-sectional images appear as discrete dots which vary in size, shape, and intensity across stacked layers along the X-axis. Because of this heterogeneity, applying a global detection threshold was not possible and required user-directed manual thresholding for each layer. To both reduce the input parameters from the user, and provide unbiased detection, cross-sectional images of the actin and nucleus channels were segmented utilizing a trained convolutional neural network based on a U-Net architecture^25, 26^.

Training and validation image sets were generated manually by labeling each F-actin dot and nuclei border on Y-Z cross-sectional images. For the training and validation sets, we randomly assigned 44 slices from two cell images. The sliced images were padded to a size of 512×512 pixels, and a total number of 44 images were split into 38 for training and 6 for validation.

Parameters for learning were: learning rate - 0.001, batch size - 1, number of epochs - 200. We used a graphics processing unit (GPU) to speed up the training process. The neural network minimizes the loss function during training that quantifies the pixel-to-pixel differences between the predicted and target image (**Fig.1b**). We changed the loss function to increase the error for false-negative results for actin fibers to 200. Apical and basal fibers were assigned based on the mean Z coordinate of each fiber point; if the mean is higher than the nucleus center, then the fiber is labeled as apical, if it is lower than the nucleus center, then the fiber is tagged as a basal fiber.

#### Reconstruction

As depicted in **Fig.1c**, to reconstruct individual F-actin structures, we grew each of the detected F-actin dots (will be referred as contours) from the first layer by connecting them to the detected contours on the next layer if it satisfied the overlap criterion from the previous layer mask. If the contour did not overlap with any other contour from the previous layer, a new actin fiber object was created. If the contour overlapped with more than one contour from the previous layer (i.e., branching points), the contour was added to a fiber whose contour had the biggest overlap area. For example, if two contours on the current layer overlapped the same contour on the previous layer, the contour with the larger overlapping area was added to the existing actin fiber object, and a new actin fiber object created for the second contour. At the end of the reconstruction, all fibers smaller than 1 µm in length were filtered out (optional parameter). Fiber length and volume were measured for each specific fiber within each cell in the dataset. Aggregated fiber statistics on a cellular level include the total fiber volume, length, and count of apical fibers, basal fibers, and the whole cell (apical + basal).

To reconstruct the nucleus (**Fig. 1d**), the contours of the nucleus shape on each Y-Z plane were combined together as a nucleus object. Volume was measured for the reconstructed object. Length was measured as the length of the rotated nucleus projection on the X-axis, width on the Y-axis, and the height extracted by applying the ellipsoid volume formula:(4/3) x π x R1 x R2 x R.

### Sample statistics and the statistical analysis of the unbiased machine learning data

The statistical analyses were based on a dataset containing 19 non-treated (-Dox) and 26 treated (+Dox) cells with 12 variables (basal fiber length, apical fiber length, total fiber length, basal fiber volume, apical fiber volume, total fiber volume, basal fiber number, apical fiber number, total fiber number, nucleus width, nucleus length, nucleus volume). To reduce the confounding bias and to obtain treated and non-treated cells with similar characteristics, propensity score matching was used with the nearest neighbor method. When calculating the propensity score, a 1:1 allocation ratio based on the nucleus volume variable was used. Thus, 19 treated cells were selected from 26 treated cells that were most statistically similar to the 19 cells in the non-treated group.

All statistical analyses were applied using R-software, version 4.1.3 (R Core Team, 2022) and the RStudio graphical interface. Shapiro-Wilk test was used to determine whether the variables were distributed normally. Continuous variables are presented as mean±standard deviation or median (quartile deviation) according to their normality (**Table 1**). The Spearman correlation coefficient was used to examine correlations between variables (**Fig.4c**). When comparing the variables between groups, bootstrap t-test with 1000 replications was preferred because of the small sample size. Two-tailed p-value <= 0.05 was considered statistically significant in analyses. In addition to p-values, r effect sizes were also calculated for comparisons.

**Table 1.**
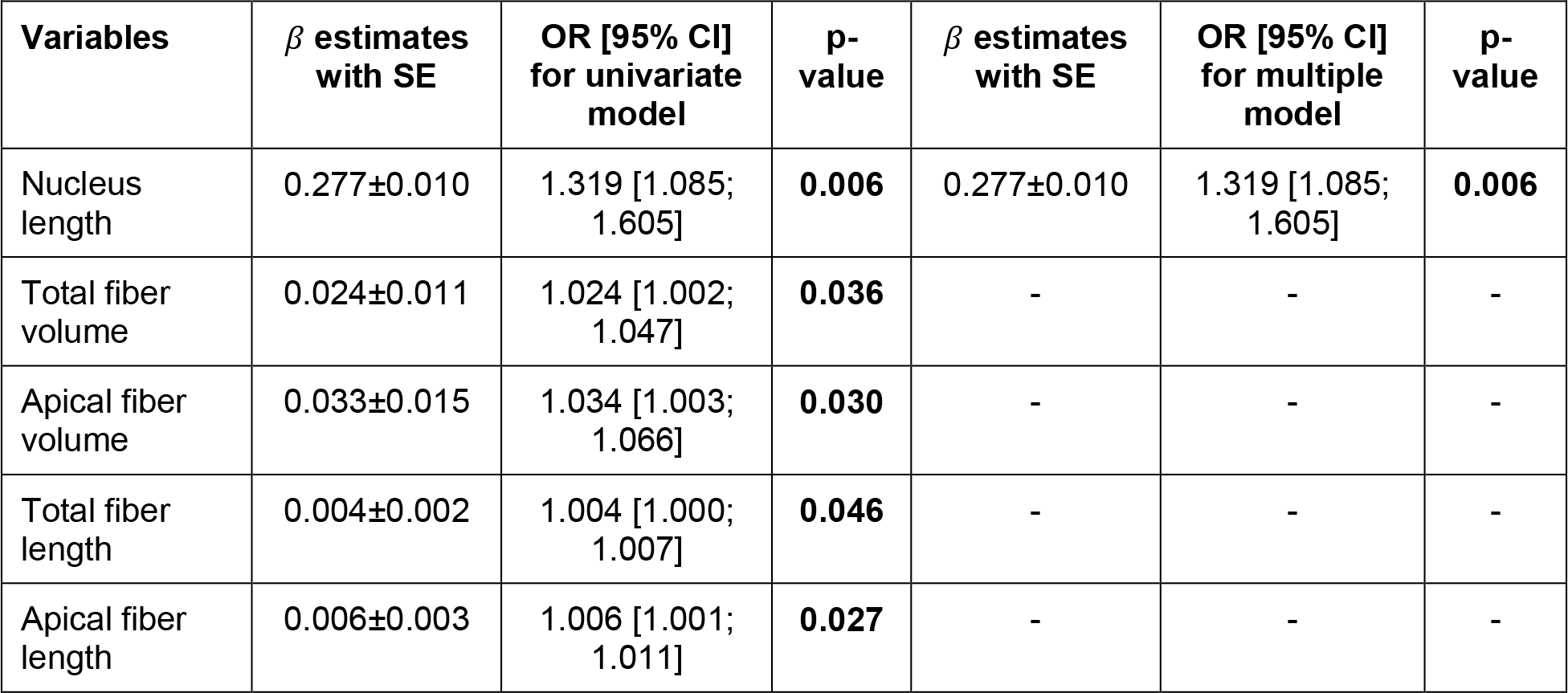
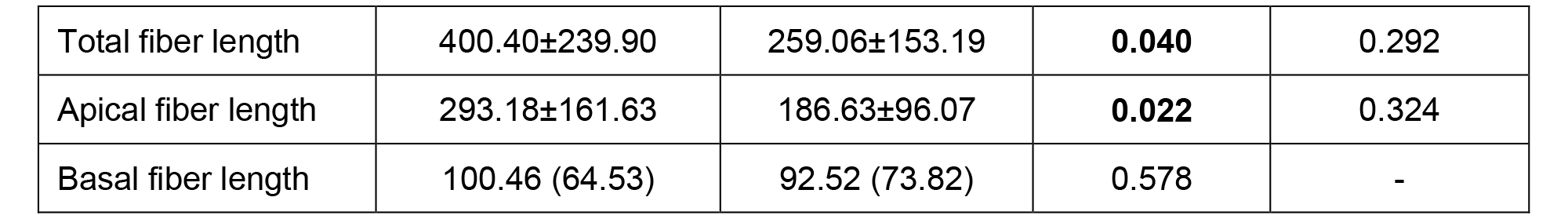
Characteristics of dataset, **bold** indicates p-value<0.005

To see the effects and odds ratios of variables both a univariate logistic regression model and a multiple logistic regression model with stepwise variable selection were used (**Table 2**). Redundant physical parameter measurements of each cell were reduced and described with fewer cell properties via a principal component analysis (PCA). Usage of PCA-transformed data, serves to prevent multicollinearity, reduces the dimension of the dataset and improves the classification performance. Number of principal components were determined via elbow criteria in the scree plot of eigenvalues (**Fig.4b**). Finally, to classify the data-points as non-treated and treated groups based on selected two principal components, three different discriminant analyses were applied: linear discriminant analysis (LDA), nonlinear quadratic discriminant analysis (QDA), and mixture discriminant analysis (MDA). Accuracy rate, sensitivity, and specificity measures of confusion matrices were used for the performance evaluation of methods. A flowchart containing the scheme for all statistical analyses is demonstrated in **Fig. S1**. Raincloud plots to visualize the summary statistics of the nucleus and fibers are added in **Fig. S2** and **Fig. S3**.

**Table 2.**
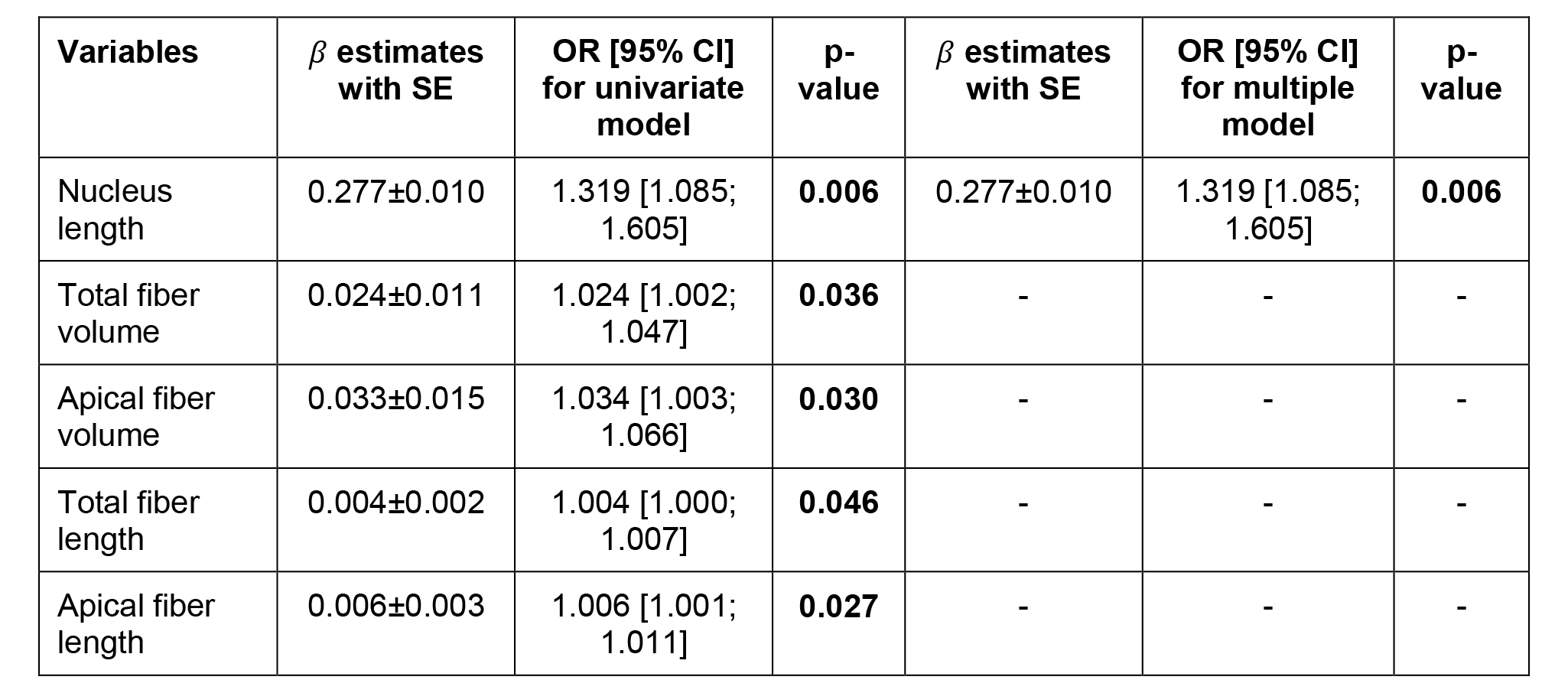
Univariate and multiple binary logistic regression results, **bold** indicates p-value<0.005

## Results

### Overexpressing Nesprin KASH domain disables LINC function in MSCs

To disable LINC function, we stably infected MSCs via lentivirus harboring a doxycycline (Dox) inducible mCherry-tagged KASH domain (dnKASH-MSCs). 1 μg/ml Dox was added to cell culture medium to induce mCherry-KASH and prevent actin linking to Nesprins on the nuclear envelope (**Fig.2a**, referred as +Dox). Controls were not exposed to Dox. Shown in **Fig.2b**, +Dox treatment increased mCherry intensity by 73% (n=515, p<0.0001) and decreased Nesprin-2 intensity by 63% (n = 530, P < 0.0001), measured over the nuclear area, indicating that Nesprin-2 was displaced from nucleus in mCherry expressing cells.

### Disabling LINC function reduces unbiased measures of apical but not basal F-actin

Shown in **Fig.3a**, +Dox treatment resulted in less F-actin fibers across the apical nuclear surface and changed both the nucleus and F-actin measures. Statistical comparison of all 12 variables between Dox treated and controls were given in **Table 1** and **Figures S2 & S3**. Total fiber volume and apical fiber volume were 67% (p=0.016) and 47% (p=0.022) smaller in the +Dox groups, respectively. Similarly, total and actin apical fiber lengths were both 37% shorter (p<0.05). As shown in and **Fig.3b**, when fibers from all cells were pooled, +Dox treatment resulted in shortened total and apical fiber lengths (15%, p<0.001). Total fiber volume and apical fiber volume were 30% (p<0.001) and 32% (p<0.001) smaller in the +Dox groups, respectively. Depicted in **Fig.3c**, length and volume distributions showed that control cells with no Dox treatment exhibited longer F-actin fibers with more volume: the longest fibers with the greatest volume in the -Dox group were 30 to 50% larger than those in the +Dox group. Basal F-actin fiber measurements did not change. Average nucleus length was also 23% smaller in the +Dox group (**Table 1**, p = 0.0007). Taken together, these results show a significant decrease in the volume and length of actin fibers associated with the apical nuclear surface by +Dox induced disruption of actin/nesprin binding, resulting in a less elongated nucleus.

### Disabling LINC function uncouples f-actin from nuclear shape measures

We next explored correlations between the 12 variables across the groups. Correlation between total fiber measures, number, length and volume, remained relatively unchanged between -Dox (0.87±0.04) and +Dox (0.94±0.03) groups. Fiber length, volume, and number all had lower correlations with the nucleus shape measures in the +Dox group when compared to the -Dox group (**Fig. 4a**). For example, the average correlation of apical F-actin number, volume and length with nuclear width, length, and volume was 0.73±0.04 in the -Dox group, dropping by 50% to 0.36±0.05 in the +Dox group. This indicated a strong decoupling between F-actin configuration and nuclear shape when LINC was disrupted due to the +Dox treatment.

**Figure 4:**
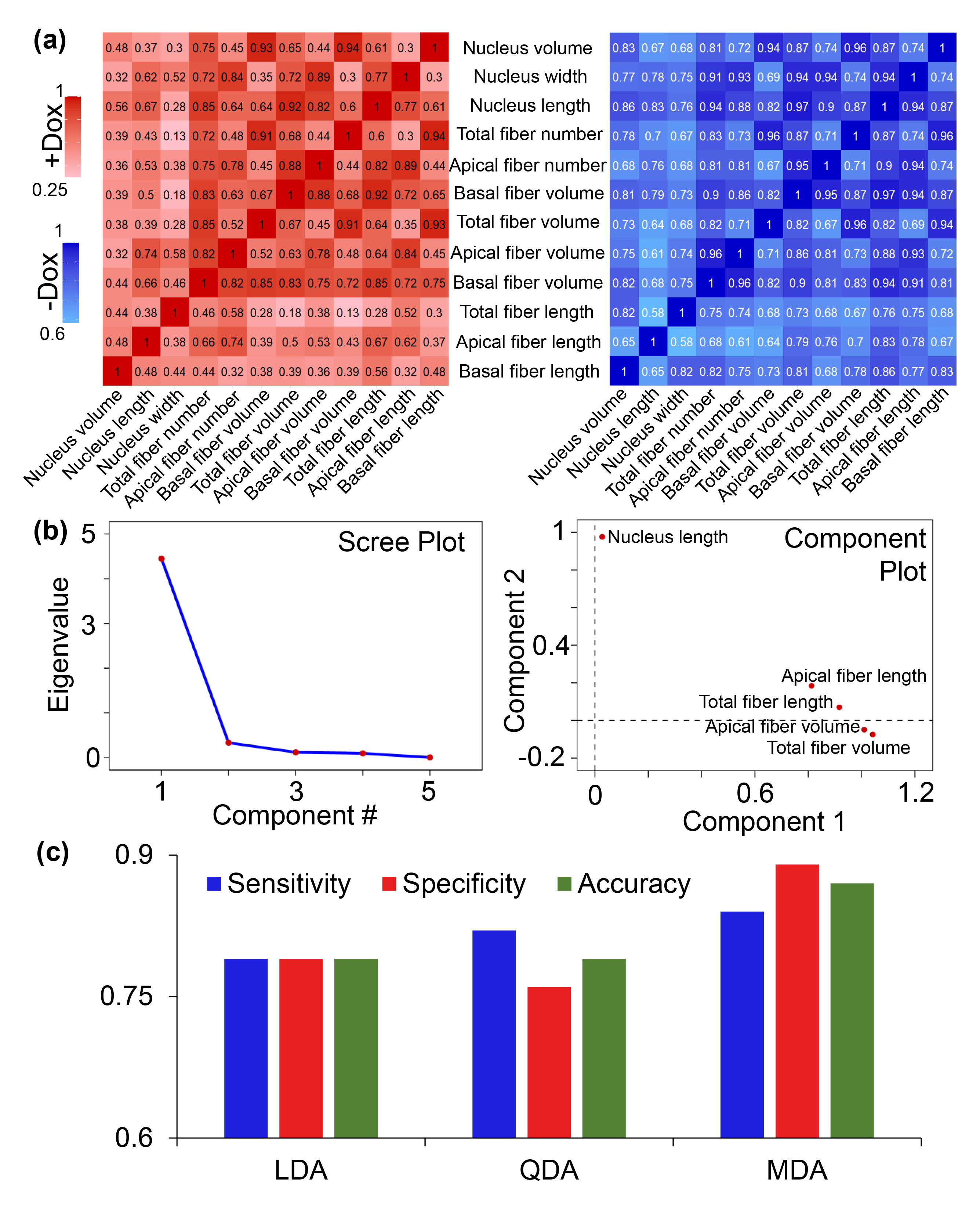
Disabling LINC function uncouples f-actin from nuclear shape measures. **(a)** Correlations of +Dox and -Dox groups, from left to right. Correlation between total fiber measures remained relatively unchanged between the +Dox (0.87±0.04) and the -Dox (0.94±0.03) groups. Average correlation of apical f-actin measures with nuclear width, length, and volume (0.73±0.04) seen a 50% drop in the +Dox group and reduced to 0.36±0.05. **(b)** Scree plot of the eigenvalues (left) and rotated principal components plot (right). Two uncorrelated principal components were found, one representing nucleus length and the others representing apical fiber length, total fiber length, apical fiber volume, and total fiber volume which explained the 95.60% of the total variance in the dataset. **(c)** Linear, quadratic and mixture discriminant analysis approaches were used to classify the cells into either treated or non-treated groups. LDA and QDA performed similarly, while MDA showed the best classification accuracy rate (87%), sensitivity (84%), and specificity (90%).

Further univariate, and multiple binary logistic regressions were applied to the dataset to find how significant variables increased the likelihood of being in one of the groups. According to **Table 2**, using the stepwise variable selection criteria, only the nucleus length variable was selected. Shown in **Table 2**, when nucleus length increases by 1 unit, the probability of being in the -Dox group increases 1.32 times compared to the +Dox, suggesting nuclear length as the most predictive measure. Next, we performed a principal component analysis (PCA). Shown in **Fig. 4b**, two uncorrelated principal components were found, one representing nucleus length and the others representing apical fiber length, total fiber length, apical fiber volume, and total fiber volume which explained the 95.60% of the total variance in the dataset. To test the accuracy of these two principal components, we performed discriminant analysis. To classify the cells into either treated or non-treated groups, quadratic and mixture discriminant analysis approaches were used. Results shown in **Fig.4c** indicate that LDA and QDA performed similarly, while MDA showed the best classification accuracy rate (87%), sensitivity (84%), and specificity (90%). Together these findings support that our unbiased approach can predict whether LINC function is disabled by querying nuclear length.

### LINC Depletion alters cell attachment and actin related gene-expression in MSCs

Finally, to understand the possible transcriptional changes due to alterations in the actin and nucleus under +Dox treatment, we performed RNA-seq analysis. DESEQ2 analyzes filtered gene pairs with significant expression differentials (p< 0.05). Shown in **Fig. 5a**, a hierarchical heatmap showed a clustering of +Dox treatments (i.e., dnKASH expression). Shown in **Fig. 5b**, a total of 177 genes (127 up, 50 down) were differentially regulated between +Dox and –Dox groups with p<0.05 statistical significance. Comparing the gene profiles between the ±Dox groups, DAVID analyses identified 38 differentially expressed pathways. Downregulated genes only associated with 2 pathways (total of 5 genes). Upregulated pathways included cell migration, integrin binding, integrin signaling and cell adhesion related pathways. Shown in **Fig.5c**, quantification of cytoskeleton and cell adhesion related genes revealed that +Dox treatment significantly increased the expression of 17 genes including Adhesion G protein-coupled receptor G1 (Agdrg1)^27^ and CD93^28^ which have roles in RhoA mediated cell spreading and migration, as well as Integrin subunit beta 3 (itgb3), integrin subunit beta 7 (itgb7) and tyrosine-protein kinase Src (Src). The DAVID pathway analyses can be found in **Tables S2** and **S3**. The increased levels of integrin and cell spreading related genes in +Dox treated cells indicate a compensatory mechanism by which cells might respond to loss of apical actin filament volume by upregulating RhoA mediated cell spreading.

**Figure 5:**
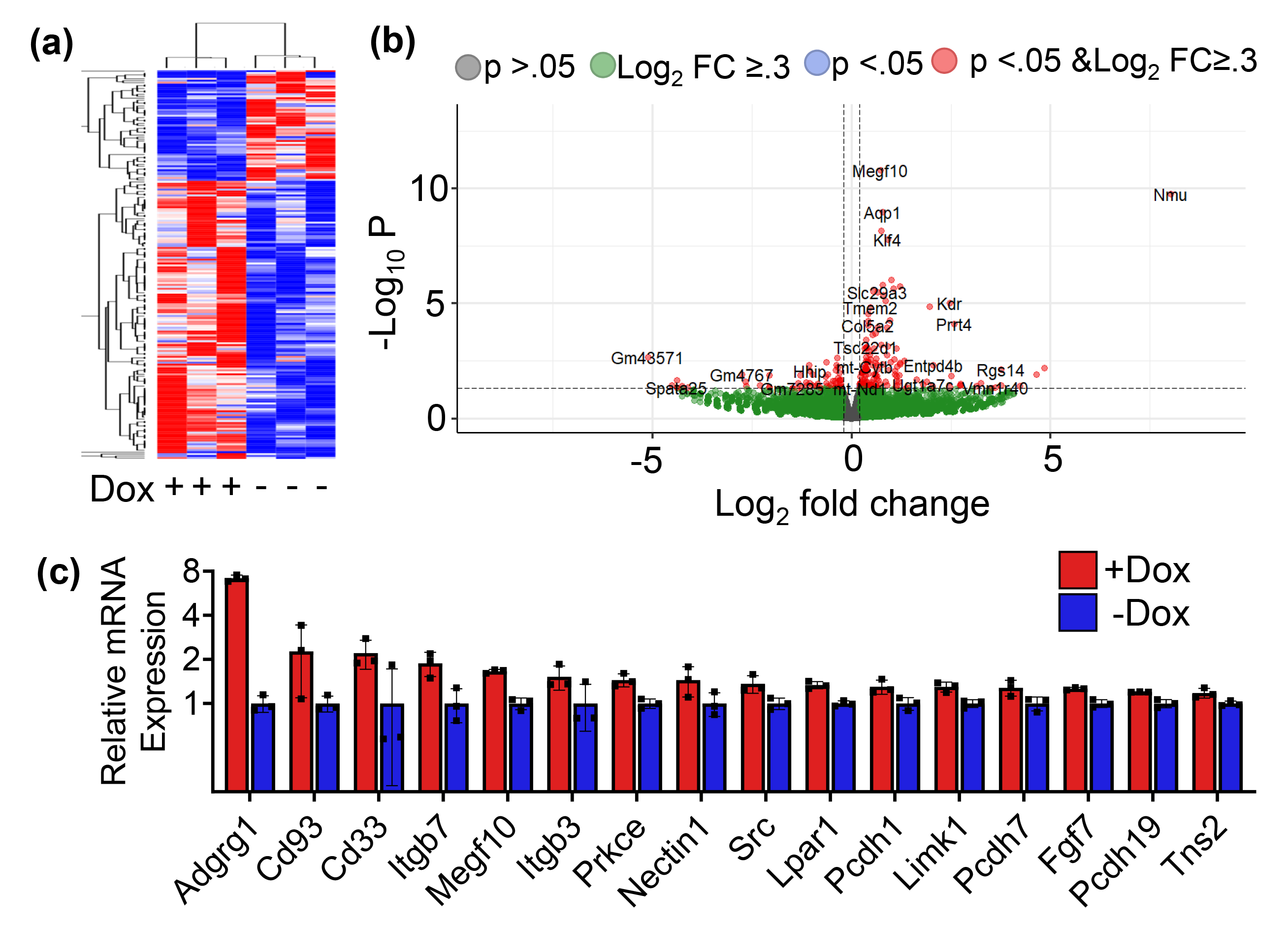
LINC Depletion alters cell attachment and actin related gene-expression in MSCs. **(a)** DESEQ2 analyzes filtered gene pairs with significant expression differentials (p< 0.05). Hierarchical heatmap showed a clustering of +Dox treatments. **(b)** Total of 177 genes (127 up, 50 down) were differentially regulated between +Dox and -Dox groups with p<0.05 statistical significance. **(c)** Quantification of cytoskeleton and cell adhesion related genes revealed significantly increased expression in +Dox treated groups.

### Conclusions and discussion

Here we developed an automated volumetric detection method for cell nuclei and nuclei-associated F-actin fibers. Our method requires no user input for segmentation and allows unbiased analysis of confocal images. Versatile post-processing options based on user needs can be adapted to allow detection of large range of F-actin structures associated with nuclei. As shown by quantification of a relatively small data set, this approach permits a comprehensive statistical analysis of F-actin structure and should lend itself to high-throughput approaches when coupled with automated data collection, available through most of the modern microscopes. As most *in vitro* investigations also rely on sampling relatively small number of cells in an imaging plate, such unbiased, high throughput capabilities that detect inherent variations in single dish will expand cytoskeletal analysis options and aid repeatability of data across multiple laboratories.

Analyzing F-actin and nuclear shape parameters in LINC disabled MSCs showed that depleting LINC function does not decrease the total number of fibers on the apical nuclear surface but instead shortens their overall length and volume. Interestingly we did not detect any changes in the number or characteristics of basal F-actin fibers, perhaps indicating that nesprin is not involved. On the apical aspect F-actin generated contractile forces indent the nuclear surface ^29–31^ magnitude of which are likely proportional to the cross-section of the F-actin fibers. Our method predicts that force across the apex of the nucleus should be reduced by disrupting LINC because F-actin volume decreased but their numbers stayed the same, likely due to a reduction in the cross section of each fiber. Such an unbiased method to detect and quantify F-actin should contribute to understand cytoskeletal forces on the nucleus. For example, mechanical models of cells often rely on simple and idealized geometries to represent cells and cytoskeleton^32–35^. Unbiased segmentation of F-actin and nuclei from confocal scans will allow more complex and realistic cellular models and thus enable researchers to quantify nuclear forces more accurately.

Using statistical models our data was able to distinguish -Dox treated cells from +Dox treated cells based on the nuclear length. This was possible because disabling LINC function reduced the correlation between F-actin and nuclear shape measures by half, indicating that F-actin regulation and nuclear shape were uncoupled in LINC disabled MSCs. Our results further indicated that the LINC disabled state was accompanied by upregulation of genes involved in cell attachment, integrin signaling and actin regulatory pathways. Similar increases in focal adhesion structure have been reported when Nesprin and Sun components of LINC complex were depleted^36–38^ or when the nucleus was softened by depleting LaminA/C^39^. To this point, we previously reported that depleting LINC function does not soften cell nuclei^40^; the preservation of nuclear modulus suggests that interfering with actin/nesprin attachment stimulates compensatory processes to losing nucleo-cytoskeletal connectivity by upregulating attachments at the cell edge.

As to effects in MSC, it has been recently reported that depleting LINC function via depletion of Sun proteins can alter heterochromatin states altering lineage selection^41^. Depleting Sun proteins results in functionally different heterochromatin rearrangements than does dnKASH expression ^21^, suggesting that changes the heterochromatin state is affected by both nuclear envelope structural composition and F-actin contractility. To this end, our method can be implemented with both fixed and live cell imaging to detect changes in F-actin under variety of mechanical forces and thus enable researchers to develop new hypotheses in cell mechanobiology.

## Abbreviations

SE: means standard error
OR: means odds ratio
CI: means confidence interval. In the models, the reference group is the treated group.

## Acknowledgements

This study was supported by AG059923, AR075803, P20GM109095, NSF1929188 and, NSF 2025505

## Data Availability

RNA-Seq data that support the findings of this study is provided as a supplementary data. The software used in this study is available on GitHub - https://github.com/mal- boisestate/afilament

## Contributions

Nina Nikitina: concept/design, data analysis/interpretation, manuscript writing

Nurbanu Bursa: data analysis/interpretation, manuscript writing

Matthew Goelzer: data analysis/interpretation, manuscript writing

Madison Goldfeldt: interpretation, manuscript writing

Chase Crandall: data analysis, final approval of manuscript

Sean Howard: data analysis, final approval of manuscript

Janet Rubin: concept/design, data analysis/interpretation, final approval of manuscript

Aykut Satici: concept/design, data analysis/interpretation, financial support, manuscript writing, final approval of manuscript

Gunes Uzer: concept/design, data analysis/interpretation, financial support, manuscript writing, final approval of manuscript

---

**Figure S1:**
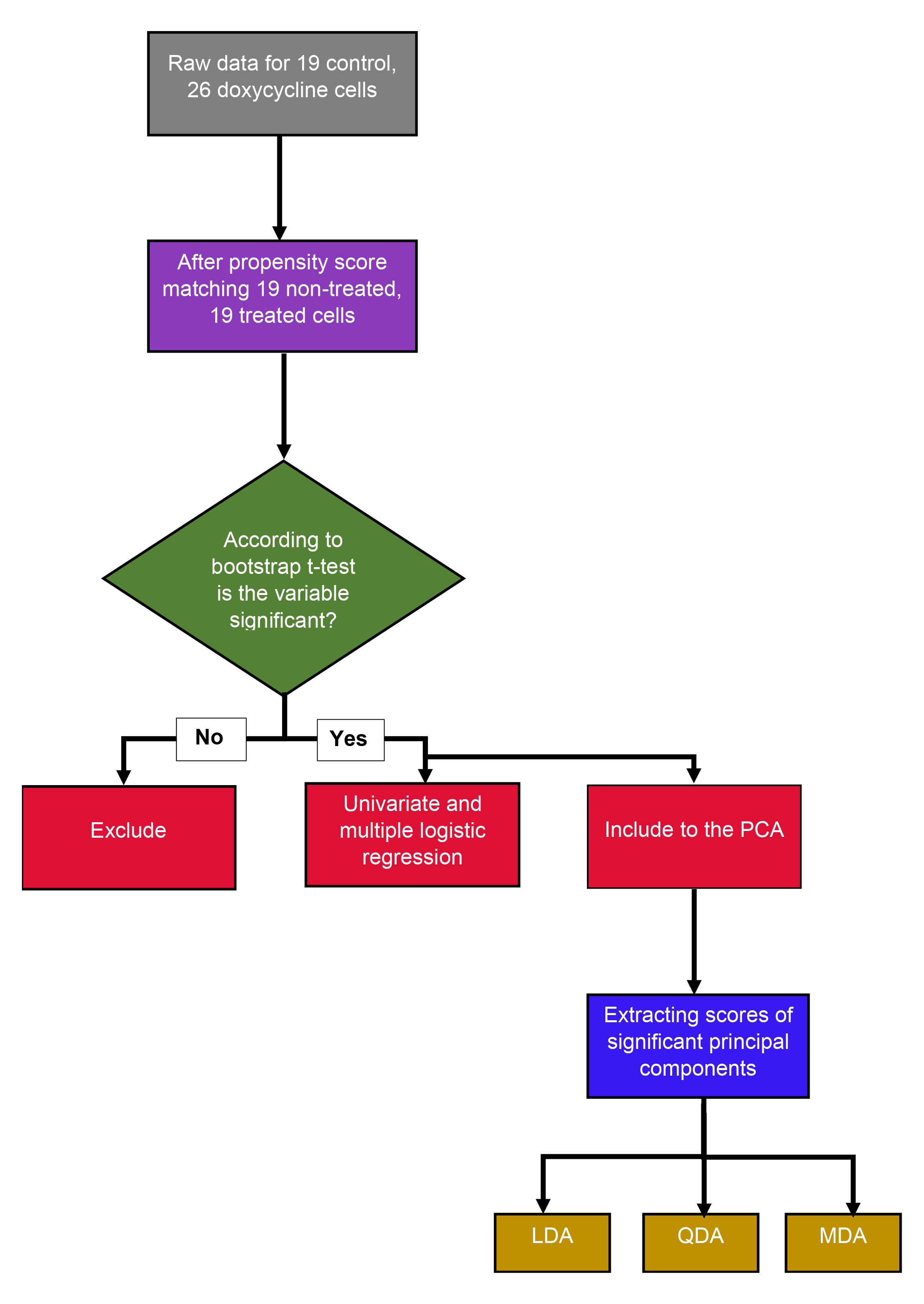
Flow chart of the statistical analyses applied to the dataset.

**Figure S2:**
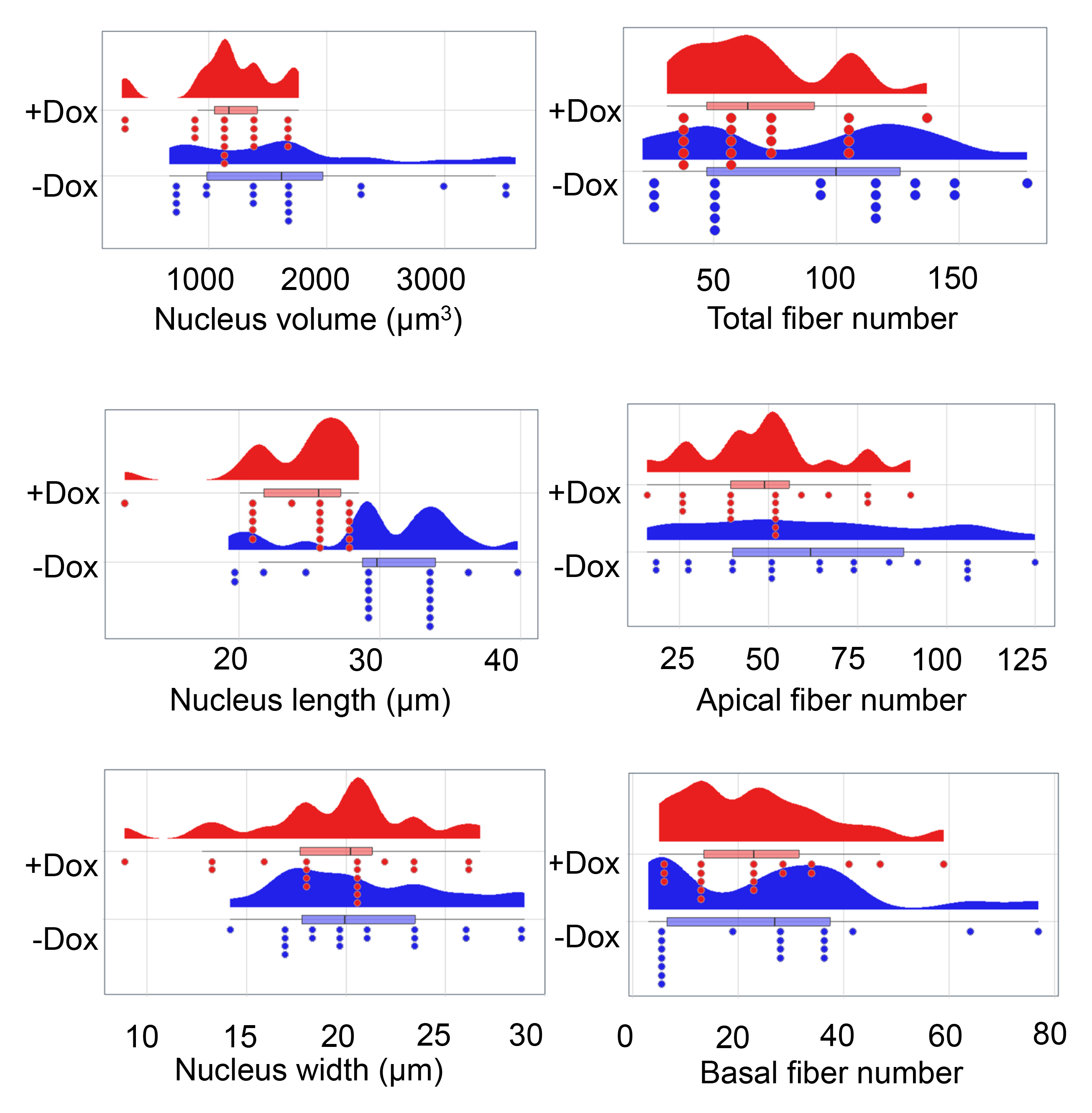
Raincloud plots for nucleus and number of fibers.

**Figure S3:**
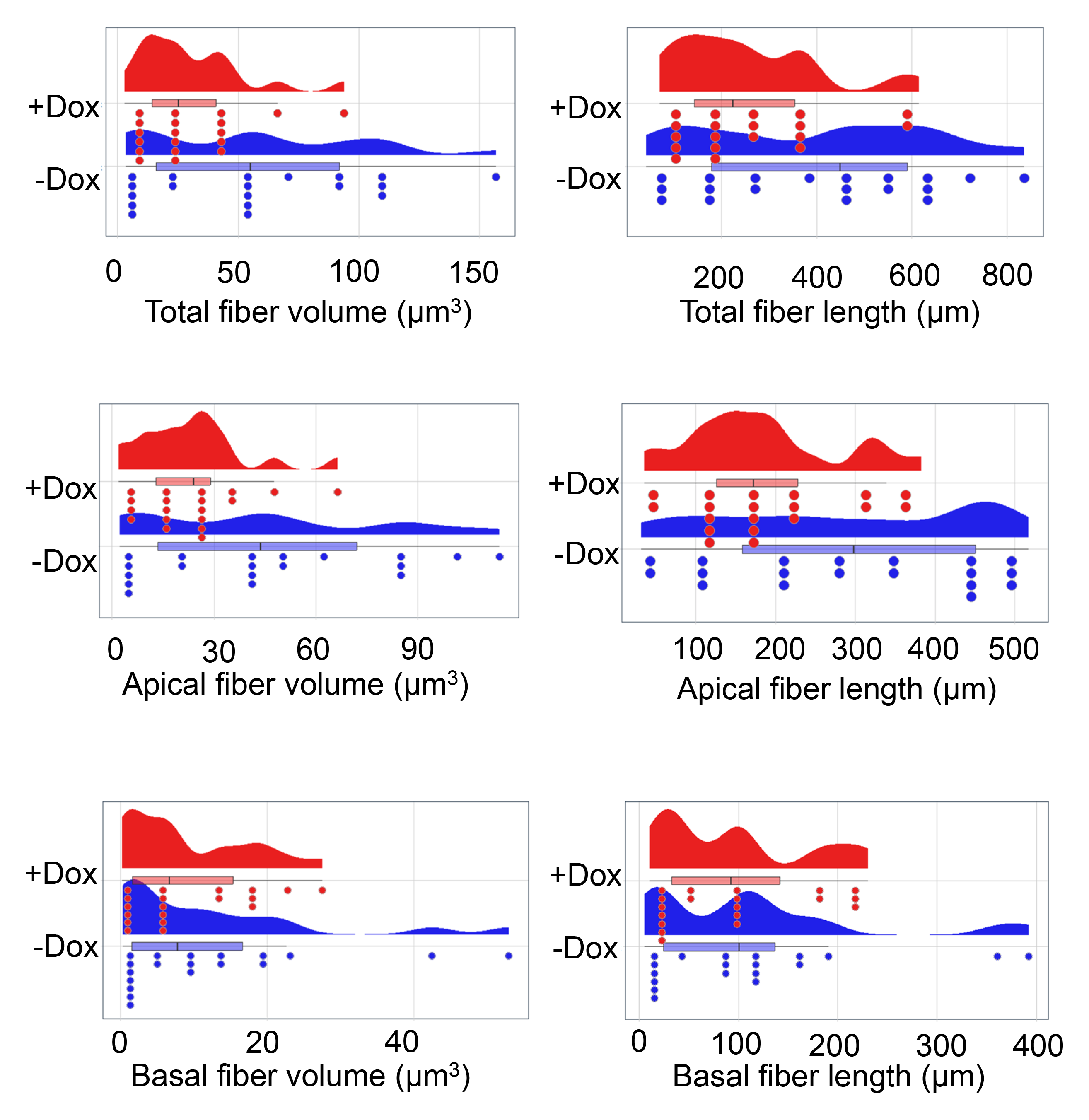
Raincloud plots for volume and length of fibers.

**Table S1:** Immunostaining antibodies and reagents and their final concentrations.

| Immunostaining antibodies and Reagents |  | Final Concentration |
| --- | --- | --- |
| Hoechst 33342 | Thermo Scientific | 1 µg/mL |
| Alexa Fluor 488 Phalloidin | Life Technologies | 0.1 µM |
| Nesprin-2 (IQ565) | Immunoquest | 1:300 |

**Table S2.**
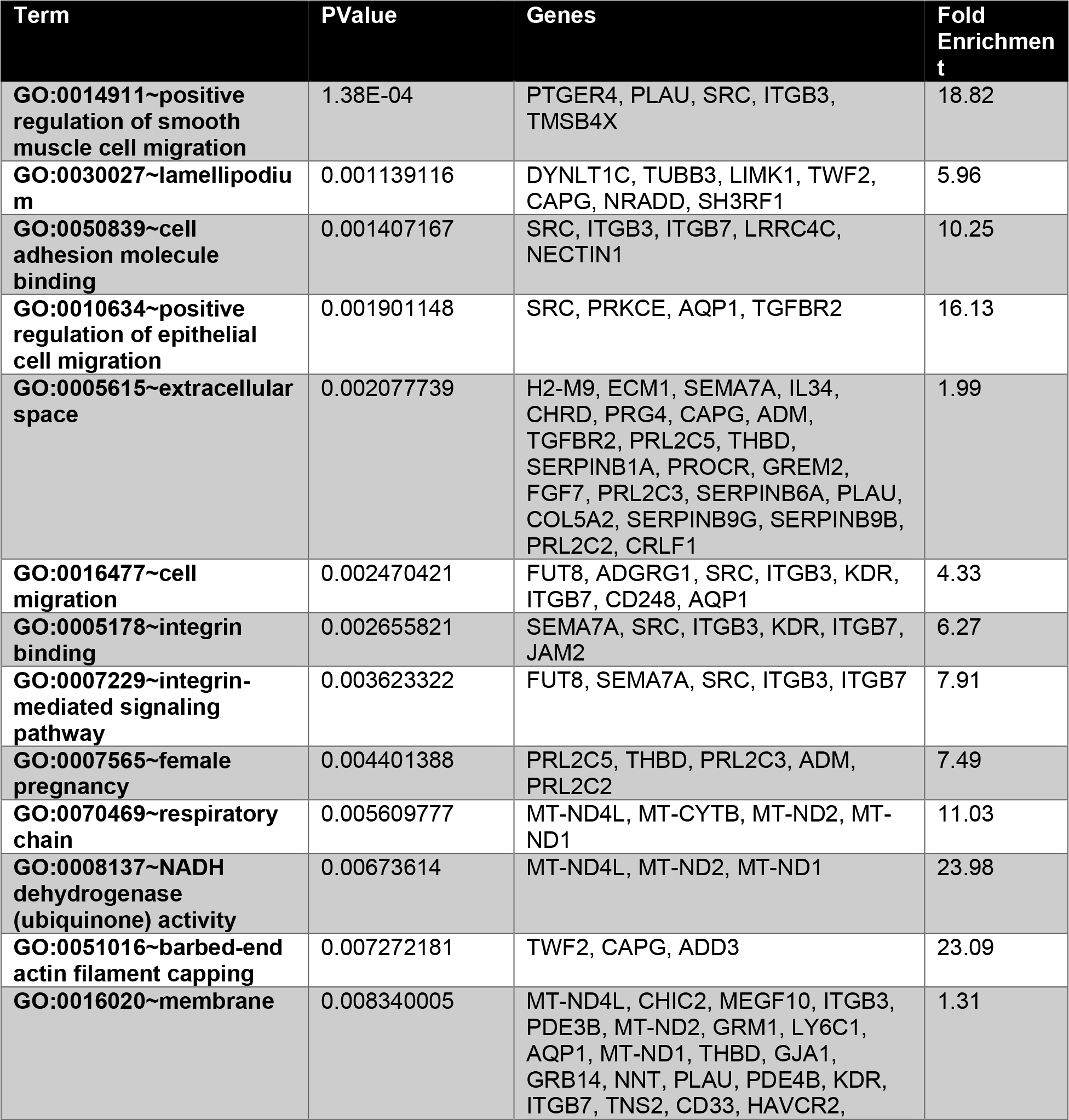
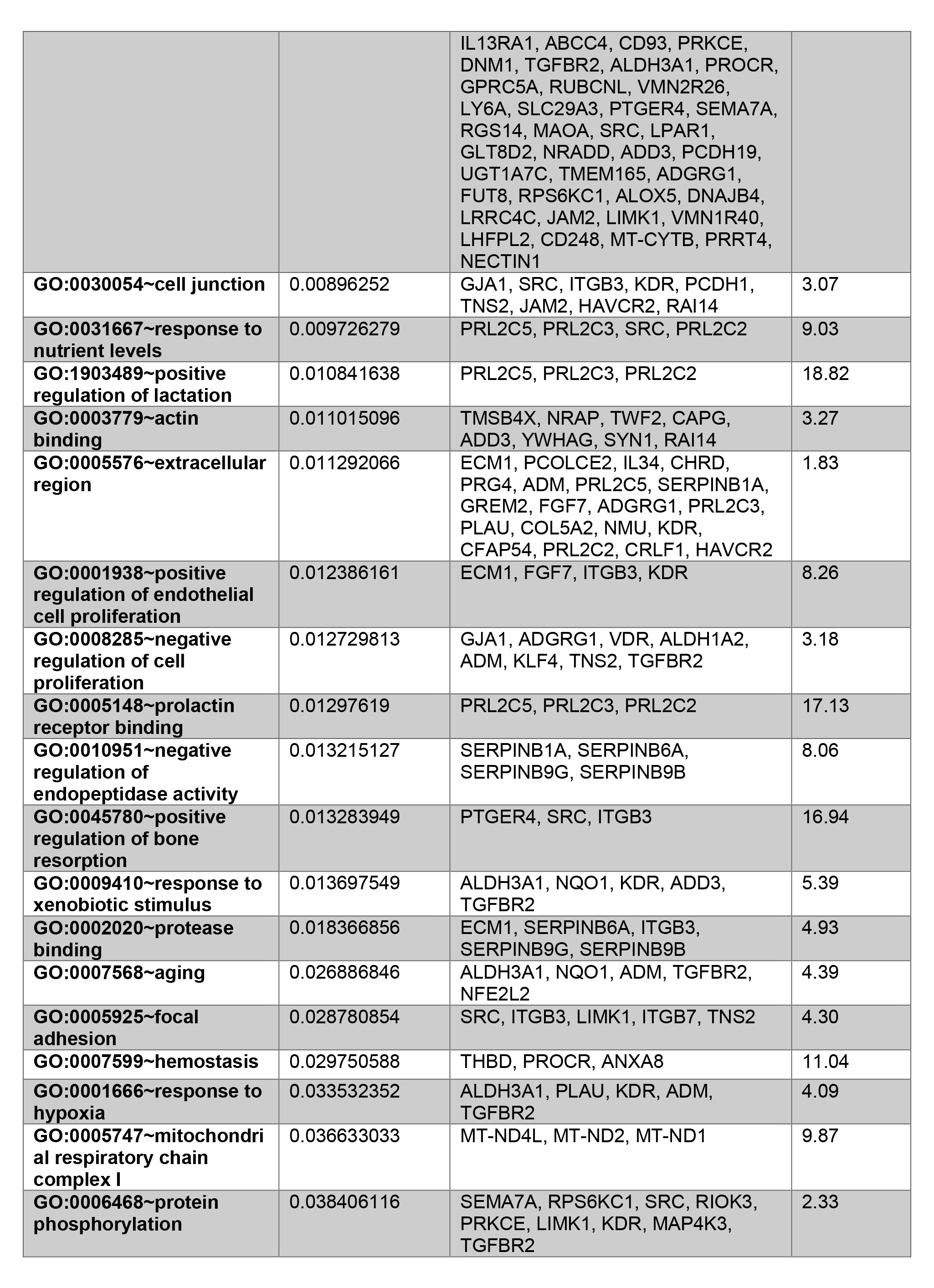
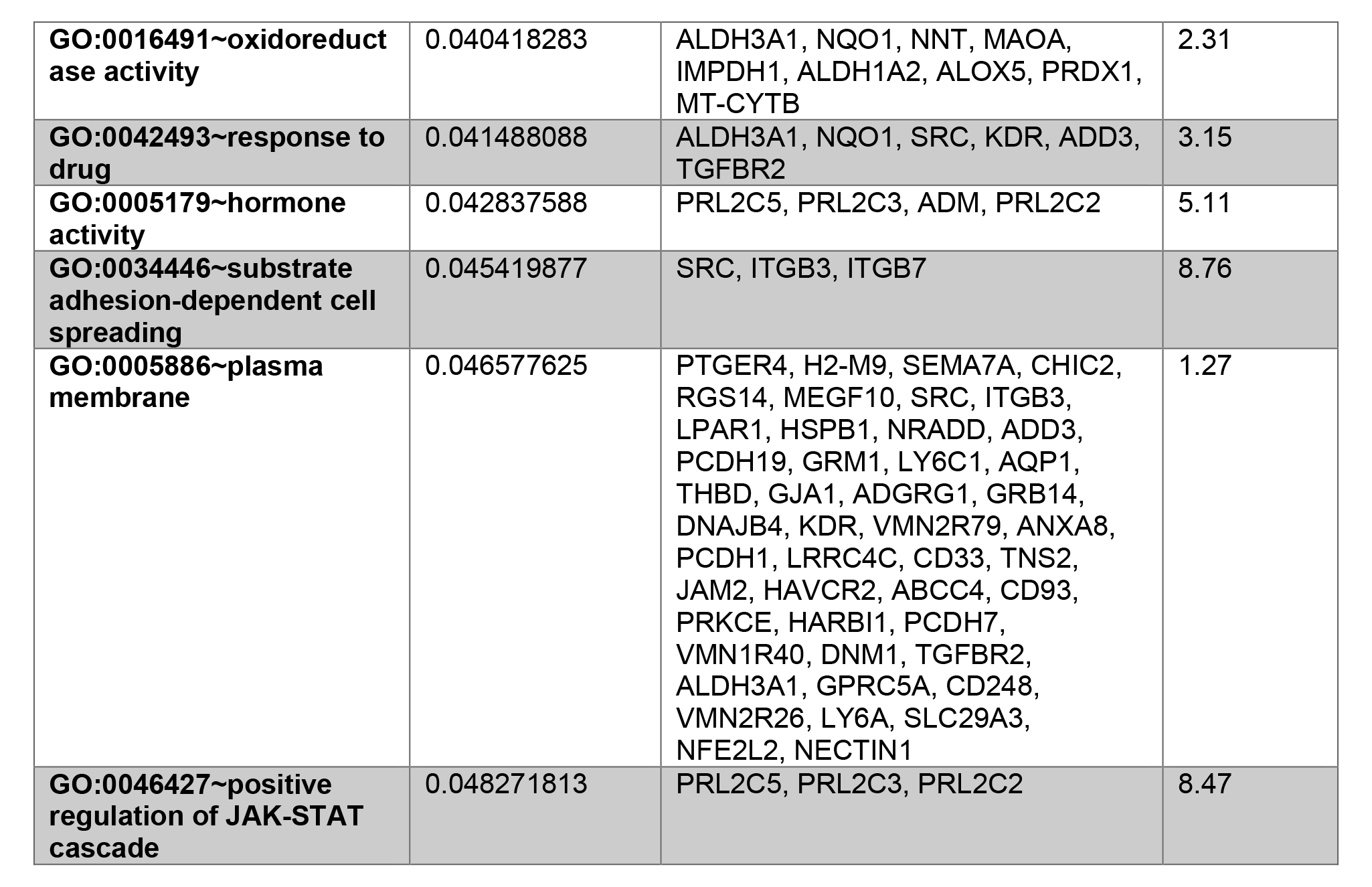
DAVID analysis of genes up regulated in +Dox vs -Dox.

**Table S3.** DAVID analysis of genes down regulated in +Dox vs -Dox.

| **Term** | **PValue** | **Genes** | **Fold Enrichment** |
| --- | --- | --- | --- |
| **GO:0008284~positive regulation of cell proliferation** | 0.018689827 | TGFB2, MARCKSL1, KLF5, GDNF, NRG1 | 4.70 |
| **GO:0008083~growth factor activity** | 0.024691651 | TGFB2, GDNF, NRG1 | 11.88 |

